# Limb regeneration modulates reproductive attributes in ladybirds: A case study in *Propylea dissecta* and *Coccinella septempunctata*

**DOI:** 10.1101/2020.08.14.251470

**Authors:** Swati Saxena, Geetanjali Mishra, Omkar

**Affiliations:** Ladybird Research Laboratory, Department of Zoology, University of Lucknow, Lucknow- 226007, India

**Keywords:** Coccinellidae, Coleoptera, forelimb, fecundity, regenerated, unregenerated

## Abstract

Regeneration is the capability to regrow or repair the lost or injured body parts. In holometabolous insects, the adult development undergoes through larval and pupal stages. Literature revealed that the limb regeneration has various impact on different life traits of organisms. In the present study, we investigated limb regeneration of two different sized ladybirds affect their life attributes. Fourth instar larvae of small ladybird *Propylea dissecta* and the large ladybird *Coccinella septempunctata* were taken from the laboratory stock and were given an ablation treatment, *viz*. forelegs of larvae were amputated at the base of the coxa. Amputated larvae were observed until the adult emergence. Emerged adults were grouped in different categories on the basis limb regeneration *i*.*e*. regenerated adults (incomplete regenerated in case of *P. dissecta*), unregenerated, and normal (control) adults. These adults were kept in different mating treatments. The unregenerated adults of both the ladybirds took more time to commence mating with shorter copulation duration and reduced fecundity and percent egg viability. Thus, it can be concluded that regeneration ability modulates the life attributes of the ladybirds irrespective of their body size.

## INTRODUCTION

Regeneration is the process of regrowth of new body structure (Shah *et al*. 2011; Gui & Yi 2002; Kumar *et al*. 2007). This phenomena is present ranging from invertebrates to vertebrates (Bely &Nyberg 2010). It is broadly classified in two categories *i*.*e*.morphallaxis where lost tissues are formed from reorganising of the existing tissue (Agata *et al*. 2007) and epimorphosis where addition of existing cells occurs. There are several tools which have been harnessed to understand the mechanism of development at genetic, cellular, tissues, organs, and organism levels (Maginnis, 2006). Mechanism of regeneration includes: (i) wound healing,(ii) blastema formation, (iii) blastema proliferation, and (iv) repatterning of de-differentiated tissue that occurs during moulting (Wigglesworth 1965). The ability of regeneration has been reported across the insect phylogeny. In holometabolous insects, such as in mulberry silkworm, *Bombyx mori* Linnaeus, ablated limbs do not regenerate as they originate from the larval prototypes (Singh *et al*. 2007). In some other holometabolous insects, development takes place in several ways. Studies on fruitfly,*Drosophila melanogaster* Meigen have revealed that the limb regeneration occurs through imaginal disc (Shah *et al*. 2011). In hemimetabolous insects, nymphs hatch out of the egg with more or less complete appendages. At genetic level, the involvement of various signalling cascades in regeneration of limbs such asJAK/STAT and Salvador/ Warts/Hippo signalling pathways (Bando *et al*. 2018) in cricket,*Gryllus bimaculatus* have been identified. Similarly, in *Tribolium castaneum* (Herbst) the role of Wnt signalling has been reported (Shah *et al*.2011). In ladybirds, the regenerated limbs have been reported in adults but not from instar to instar (Wu *et al*.2015; Saxena *et al*. 2016).

Though regeneration is beneficial to insects in terms of physical fitness, it also modulates the life attributes. Males have been known to regulate the courtship, copulation duration, quality and quantity of ejaculates depending upon the surrounding environment and their physical fitness (Wedell *et al*. 2002; Ortigosa and Rowe, 2002; Wilder and Rypstra, 2007). Poorly regenerated or physically disabled males were known to affect the mobility, mating success and reproduction (Juanes and Smith, 1995). Besides this, it has been reported that the missing or regenerated limb are known to affect the outcome of the various ecological interactions such as prey-predator interactions and intraspecific competition (O’Neill and Cobb, 1979; Garvey *et al*.,1994). In wolf spider, *Schizocosa ocreata* it has been reported that their prey capturing efficiency decreased due to missing or regenerating limb (Wrinn and Uretz, 2008). In field cricket *Gryllus bimaculatus* it has been reported that in females, loss limb has resulted in significantly reduced mating ability while in males it has resulted in reduced longevity. Under promiscuous conditions, poor quality males were overpowered by the healthy males (Schneider *et al*. 2000; Elgar *et al*., 2003). Females generally tend to invest in those males which can gain them direct or indirect benefits (Andersson, 1994), since they expect to pass down the healthier alleles to their offspring (Zahavi 1975; Iwasa *et al*. 1991).

In terms of reproductive performance, in spiders, it has been also reported that loss of forelegs can lead to the reduced reproductive success by less transfer of sperms (Johnson and Jakob, 1999). Studies in *Harmonia axyridis* have also reported that the unregenerated adults were poor performers in terms of mating and reproductive parameters than the regenerated and non-ablated individuals (Wang *et al*.2015). Ladybirds are polyandrous (Majerus 1994; Colares *et al*. 2015) and display mate choice (Kearns *et al*. 1992; Mishra and Omkar, 2014). In *Menochilus sexmaculatus* Fabricius it has been proven earlier that the regeneration occurs in adult stage and not from instar to instar and unregenerated adults were poor performers in terms of mating and reproductive parameters (Saxena *et al*.2016). Another study on *M. sexmaculatus* has reported when the adult legs were ablated from three different joints then there was the difference in their mating performance and the reproductive output (Shandilya *et al*.2018).

In the present study we aim to study the effects of limb regeneration on two ladybirds *i*.*e. Coccinella septempunctata* (L.) and *Propylea dissecta* (Mulsant). *Coccinella septempunctata* commonly known as seven spotted ladybird is a large sized species of ladybirds. They are euryphagous in nature as larvae feed on aphids to complete their life cycle while adults can feed on non-aphid prey also (Richards & Evans, 1998). *Propylea dissecta* is an Asian ladybird. They are polymorphic in nature and three morphs have been reported *i*.*e*. pale, intermediate, and typical (Pervez, 2002). In this study we have hypothesised that regeneration will have some costs in terms of reproductive attributes. For this, we took fourth instar of *P. dissecta* and *C. septempunctata* and amputated their forelimb from the base of coxa. All the adults (regenerated, normal and unregenerated) will be kept in different mating treatments and its impact on mating and reproductive attributes of both the beetles were subsequently observed.

## MATERIALS AND METHODS

### Stock maintenance

Two species of ladybirds, *i*.*e*. a medium sized beetle, *Propylea dissecta* (13.04±0.15 mg) and large beetle, *Coccinella septempunctata* (21.70±0.15 mg) (n=50 each species) were collected from the agricultural fields of Lucknow, India (26°50′N, 80°54′E). These beetles were selected for experimentation due to their predominance in local fields, wide prey range (Agarwala & Yasuda 2000) and high reproductive output. Males and females were paired in plastic Petri dishes (hereafter, 9.0×2.0 cm) and provided with *ad libitum* supply of cowpea aphid, *Aphis craccivora* Koch (Hemiptera: Aphididae) raised on cowpea, *Vigna unguiculata* L.(reared in a glasshouse at 25 ±2°C, 65 ±5% Relative Humidity). Petri dishes with mating pairswere placed in Biochemical Oxygen Demand (BOD) incubators (Yorco Super Deluxe, YSI-440, New Delhi, India) at 27 ± 1°C, 65 ±5% R.H., 14L:10D. They were inspected twice daily (1000 and 1500 h) for oviposition. The eggs were separated and reared individually in Petri dishes until the emergence of fourth instar.

### Limb amputation

Fourth instar larvae (24h old post moulting) of *P. dissecta and C. septempunctata*, were taken from the stock and divided into two groups (of 100 individuals each). First group was reared as control (termed as normal hereafter) and chilled for 5 minutes with no amputation treatment. The second group of larvae were also chilled for 5 minutes to ease the ablation process. Amputation of fore limb from base of coxa of right side was done under a stereoscopic binocular microscope (Magnus) at 16x magnification with the help of micro-scalpel. Post amputation, larvae were reared individually in Petri dishes until adult emergence. All the adults were isolated and reared on the *ad libitum* supply of *A. craccivora*.

### Mating combinations and attributes

10-day old unmated adults of both the species were taken from the amputation treatment and were assessed for limb regeneration. The ones with limb regeneration were called regenerated (in *P. dissecta*, limbs were incompletely regenerated so here we used incompletely regenerated adults) and those without regeneration were called unregenerated. These adults and normal adults were allowed to mate in following combinations (1)Regenerated♂×Regenerated♀(♂_R_× ♀_R_), (2) Regenerated♂ × Normal♀ (♂_R_× ♀_N_), (3)Regenerated♂×Unregenerated♀(♂_R_×♀_U_), (4)Unregenerated♂× Unregenerated♀ (♂_U_× ♀_U_), (5) Unregenerated♂ × Normal♀(♂_U_× ♀_N_), (6) Unregenerated♂ × Regenerated♀ (♂_U_× ♀_R_), (7) Normal♂ × Normal♀ (♂_N_× ♀_N_), (8) Normal♂ × Unregenerated♀(♂_N_× ♀_U_), and (9) Normal♂ × Regenerated♀ (♂_N_× ♀R). In case of *P. dissecta*, complete regeneration was not observed. Therefore, incompletely regenerated adults were used for forming above mating combinations. Pairs were allowed to mate until they disengaged naturally. Time to commence mating (introduction in Petri dish to establishment of genital contact) and copulation duration (from establishment of genital contact till natural disengagement) were recorded. Females were separated and kept individually in Petri dishes with *ad libitum* supply of *A. craccivora*. They were inspected twice (10:00 and 15:00hr) daily for oviposition for next seven days and the egg hatching was recorded. All the mating combinations were replicated 15 times.

### Statistical Analysis

Data on mating (time of commencement of mating and copulation duration) and reproductive attributes (fecundity and percent egg viability (dependent factors) were initially tested for normal distribution (Kolmogorov-Smirnoff test). On being found normally distributed with homogeneous variation, data on mating and reproductive attributes were subjected to two-way analysis of variance (ANOVA) with regeneration status of male and female as independent factors. This analysis was followed by comparison of means using post hoc Tukey’s honest significance test at 5%. All statistical analyses were conducted using R studio Version 1.2.1335 statistical software. The correlation analysis between copulation duration and reproductive parameters of all mating treatments of both the ladybirds were subjected to linear regression analysis using SPSS Statistic 20 software.

## RESULTS

In *P. dissecta*, regeneration status of males and females were found to have significant effect on the time to commence mating (TCM) (F_♂_=7.05, P<0.05, df=2,126; F_♀_=27.88, P<0.05, df=2, 126). The interaction between the two independent factors was insignificant (F_♂×♀_=1.89; P>0.05; df=4,126). A significant effect of regeneration status of male and female was also observed on time of commencement of mating in *C. septempunctata* (F_♂_=48.43; P<0.05; df=2,126; F_♀_=65.51; P<0.05; df=2, 126). The interaction between the two factors was also found significant (F_♂×♀_=7.69; P<0.05; df=4,126). The highest time to commence mating was found in *P. dissecta* and *C. septempunctata* when unregenerated males were allowed to mate with unregenerated females (Fig. 1a). However, in case of *P*.*dissecta*, lowest time to commence mating was observed when normal males mated with normal females, which was contrary to *C. septempunctata* where lowest TCM was observed when regenerated adults were allowed to mate (Fig. 1a).

**Figure 1:**
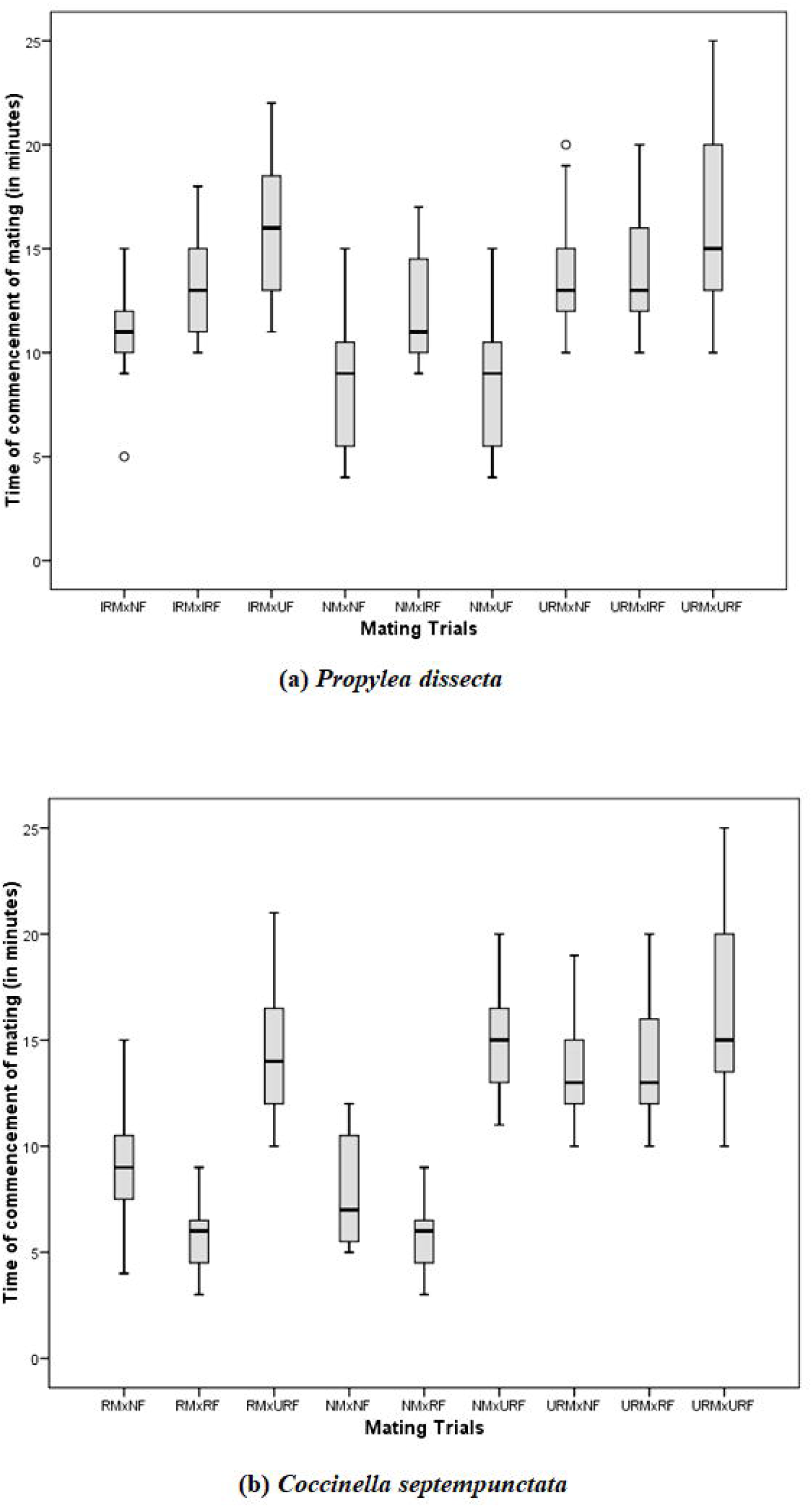
Box and whisker plots showing time of commencement of mating of (a) *Propylea dissecta* (b) *Coccinella septempunctata* when kept under dfferent mating trials. The horizontal lines lines within the box marks the median. The vertical lines extending from box are 15 times length of the box. Circles represent outliers. IRM=Incomplete regenerated males; IRF=Incomplete regenerated females; RF=Regenerated females; RM=Regenerated males; NF=Normal females; NM=Normal males; UF=Unregenerated females; UM=Unregenerated males.

Copulation duration (CD) was also found to have a significant effect of regeneration status of males and females in *P*.*dissecta* (F_♂_=175.55; P<0.05; df=2,126; F_♀_=195.35; P<0.05; df=2, 126) as well as in *C. septempunctata* (F_♂_=343.90; P<0.05; df=2,126; F_♀_=333.08; P<0.05; df=2, 126). The interactions between status of males and females were also found significant in *P*.*dissecta* (F_♂×♀_=36.12; P<0.05; df=4,126) and *C. septempunctata* (F_♂×♀_=74.36; P<0.05; df=4,126). The shortest copulation duration was recorded in unregenerated treatments (Figure 2b). In *P. dissecta*, longest copulation duration was found when normal males were paired with normal females, while in *C. septempunctata* the maximum copulation duration was recorded when regenerated adults were paired (Fig. 2b).

**Figure 2:**
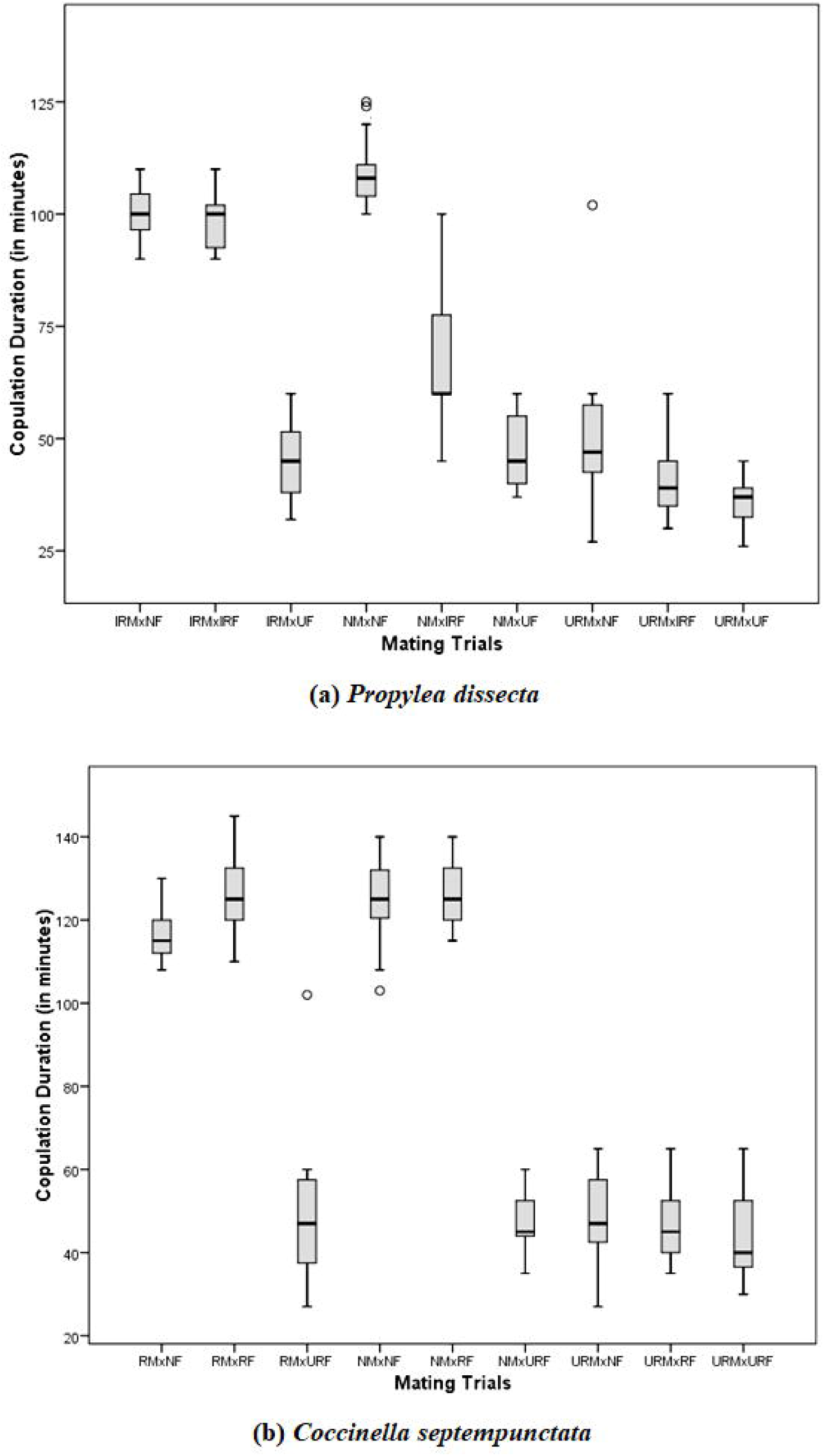
Box and whisker plots showing copulation duration of (a) *Propylea dissecta* (b) *Coccinella septempunctata* when kept under different mating trials. The horizontal lines lines within the box marks the median. The vertical lines extending from box are 15 times length of the box. Circles represent outliers. IRM=Incomplete regenerated males; IRF=Incomplete regenerated females; RF=Regenerated females; RM=Regenerated males; NF=Normal females; NM=Normal males; UF= Unregenerated females; UM:=Unregenerated males.

### Reproductive attributes

Fecundity was significantly influenced by the regeneration status of males and females in *P. dissecta* (F_♂=_1324.81; P<0.05; df=2,126; F_♀_=42.79; P<0.05; df=2, 126) and *C. septempunctata* (F_♂=_763.5; P<0.05; df=2,126; F_♀_=1018.7; P<0.05; df=2, 126). The interactions were also significant in both *P. dissecta* (F_♂×♀_=4.98; P<0.05; df=4,126) and *C. septempunctata* (F_♂×♀_=299.2; P<0.05; df=4,126). In *P*.*dissecta*, maximum fecundity was recorded in normal adults while in *C. septempunctata* it was found in regenerated adults (Fig. 3a and 3b). In both the ladybirds minimum fecundity was for unregenerated pairs (Fig. 3a and 3b). A significant effect of regeneration status of males and females was also recorded for the percent egg viability of *P*.*dissecta* (F_♂=_1051.78; P<0.05; df=2,126; F_♀_=10.42; P<0.05; df=2, 1326) and *C*.*septempunctata*(F_♂=_126.32; P<0.05; df=2,126; F_♀_=22.07; P<0.05; df=2, 126). The interaction of these factors were also significant in *P. dissecta* (F_♂×♀_=8.72; P<0.05; df=4,126) but insignificant in *C. septempunctata* (F_♂×♀_=2.02; P>0.05; df=4,126). Minimum percent egg viability was recorded for unregenerated treatments in both the ladybird beetles while maximum egg viability was found for normal treatments in *P*.*dissecta* and in regenerated treatments for *C. septempunctata* (Fig. 4a and 4b).

**Figure 3:**
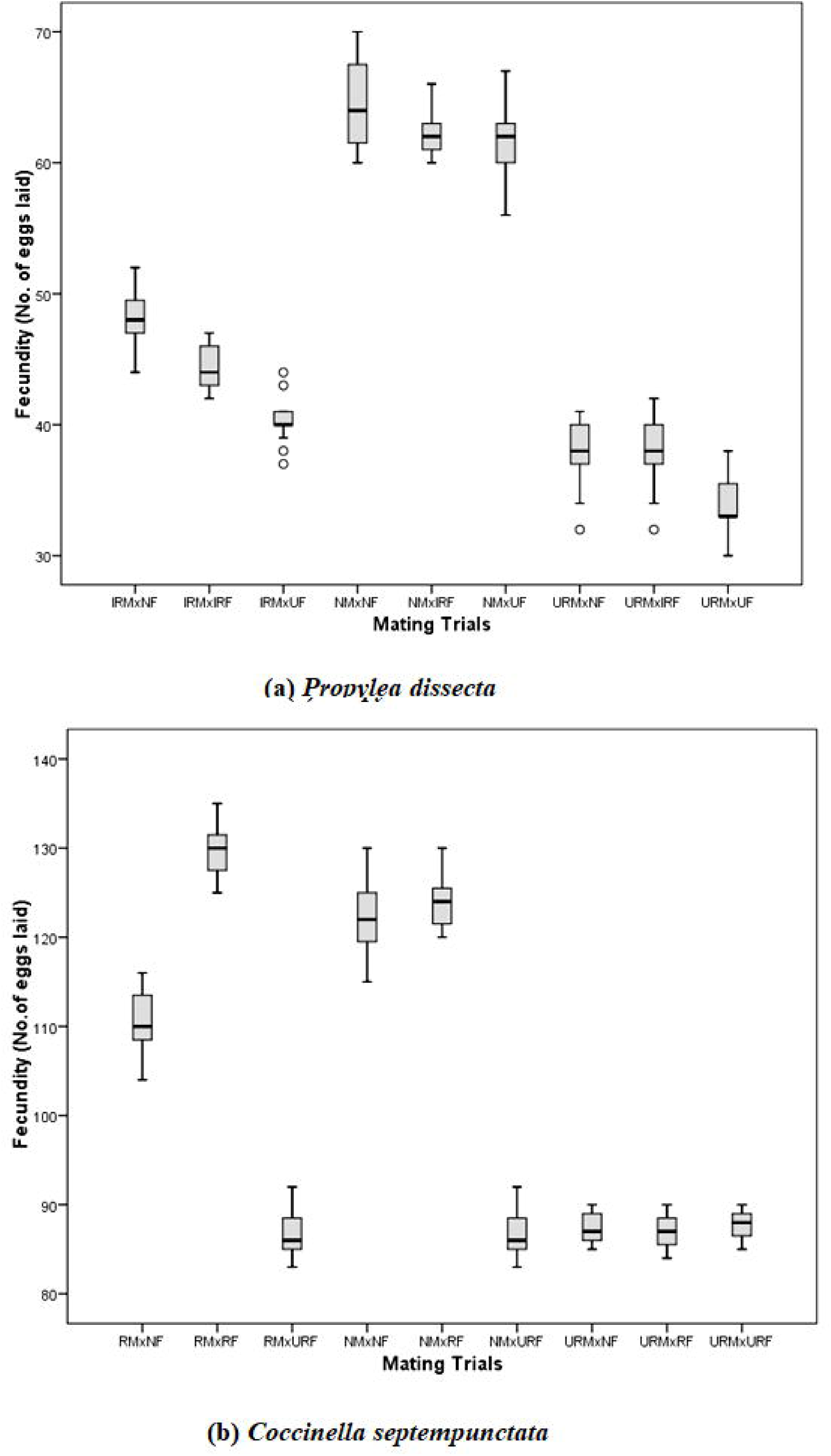
Box and whisker plots showing feundity of (a) *Propylea dissecta* (b) *Coccinella septempunctata* when kept under different mating trials. The horizontal lines lines within the box marks the median. The vertical lines extending from box are 15 times length of the box. Circles represent outliers. IRM=Incomplete regenerated males; IRF=Incomplete regenerated females; RF=Regenerated females; RM=Regenerated males; NF=Normal females; NM=Normal males; UF=Unregenerated females; UM=Unregenerated males.

**Figure 4:**
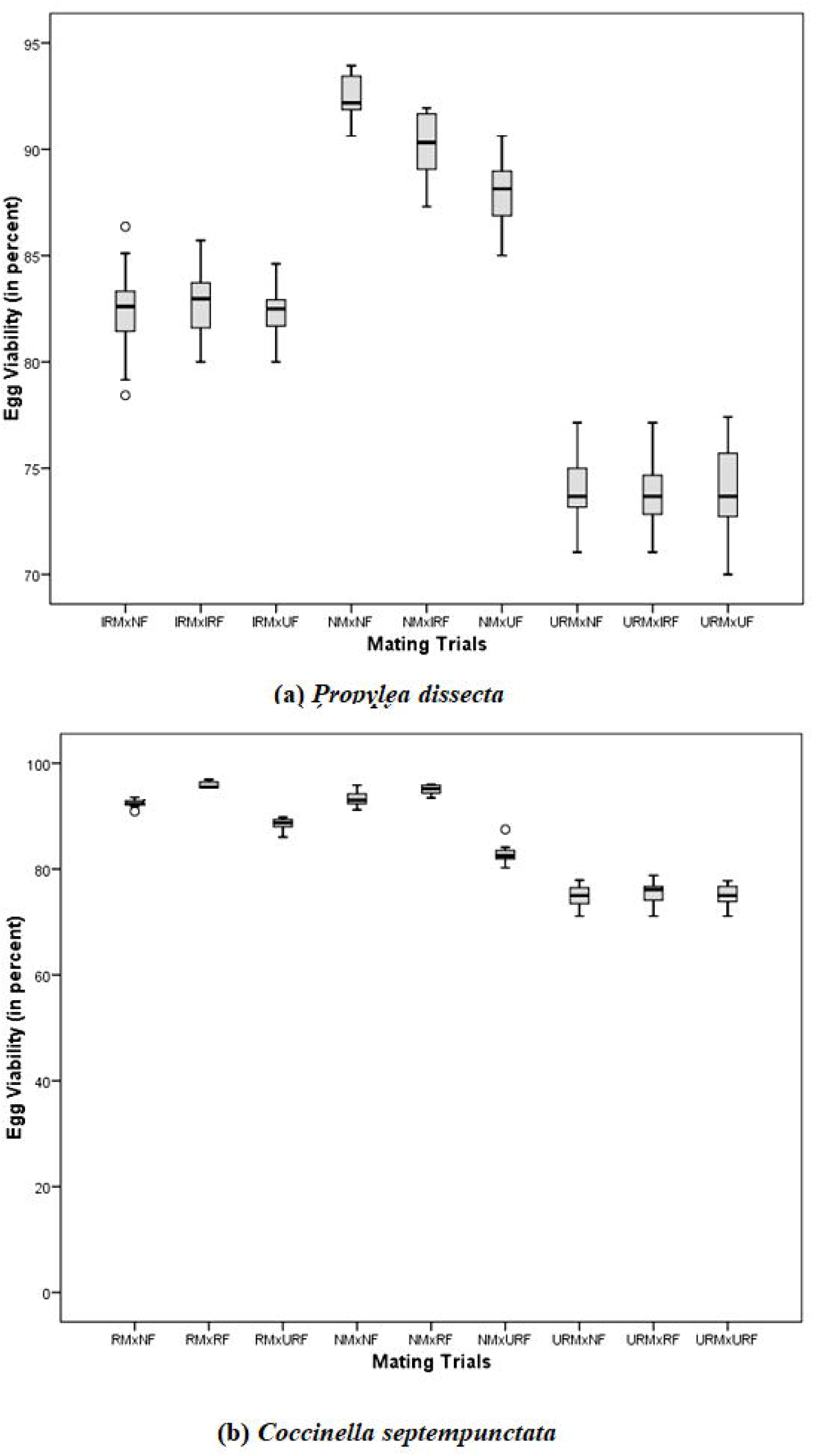
Box and whisker plots showing percent viability of (a) *Propylea dissecta* (b) *Coccinella septempunctata* when kept under different mating trials. The horizontal lines lines within the box marks the median. The vertical lines extending from box are 15 times length of the box. Circles represent outliers. IRM=Incomplete regenerated males; IRF=Incomplete regenerated females; RF=Regenerated females; RM=Regenerated males; NF=Normal females; NM=Normal males; UF=Unregenerated females; UM=Unregenerated males.

**Figure 5:**
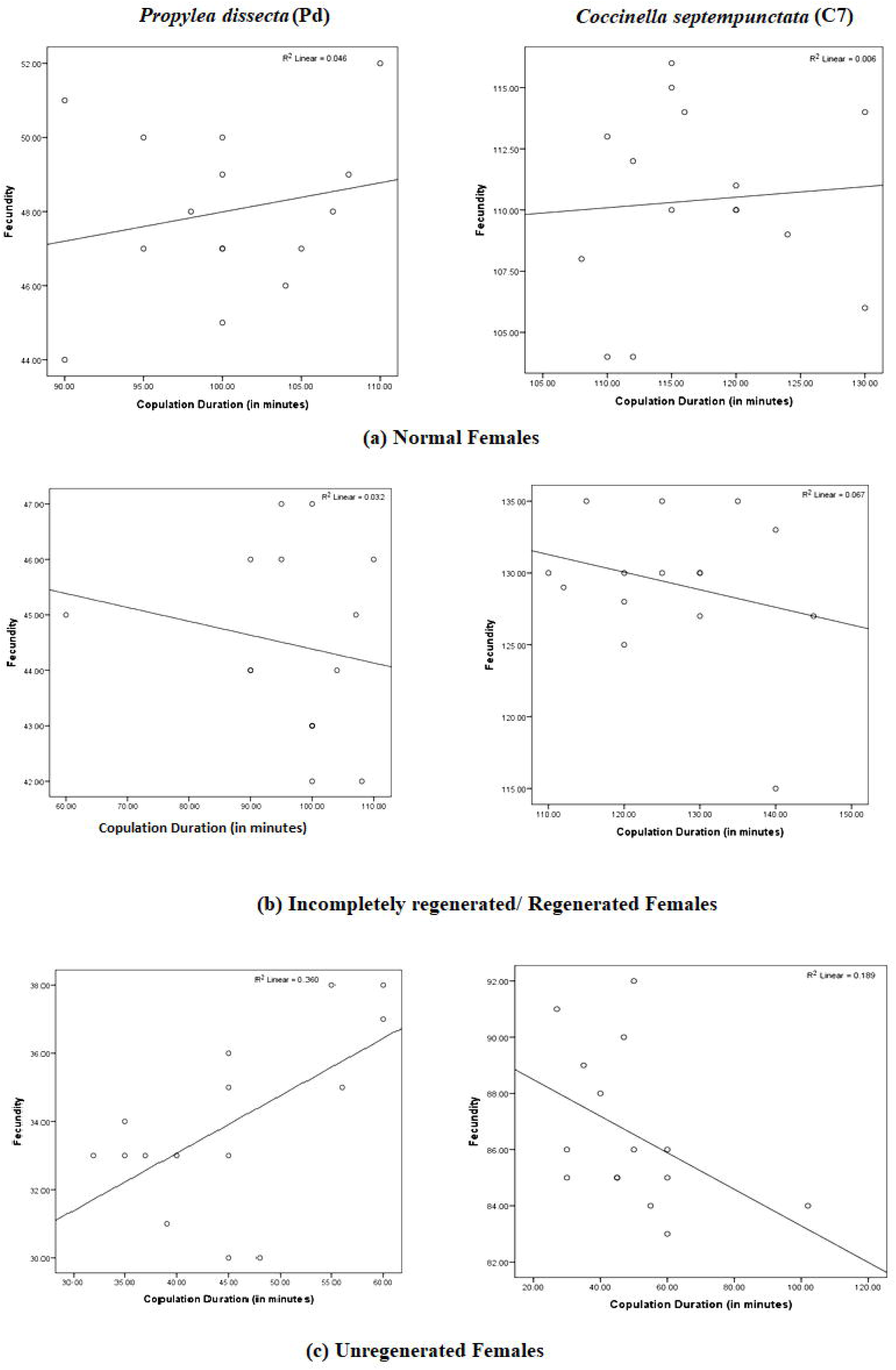
Relation between copulation duration and fecundity when different females of Pd and C7 mated with regenerated (incompletely in *Propylea dissecta)* males.

**Figure 6:**
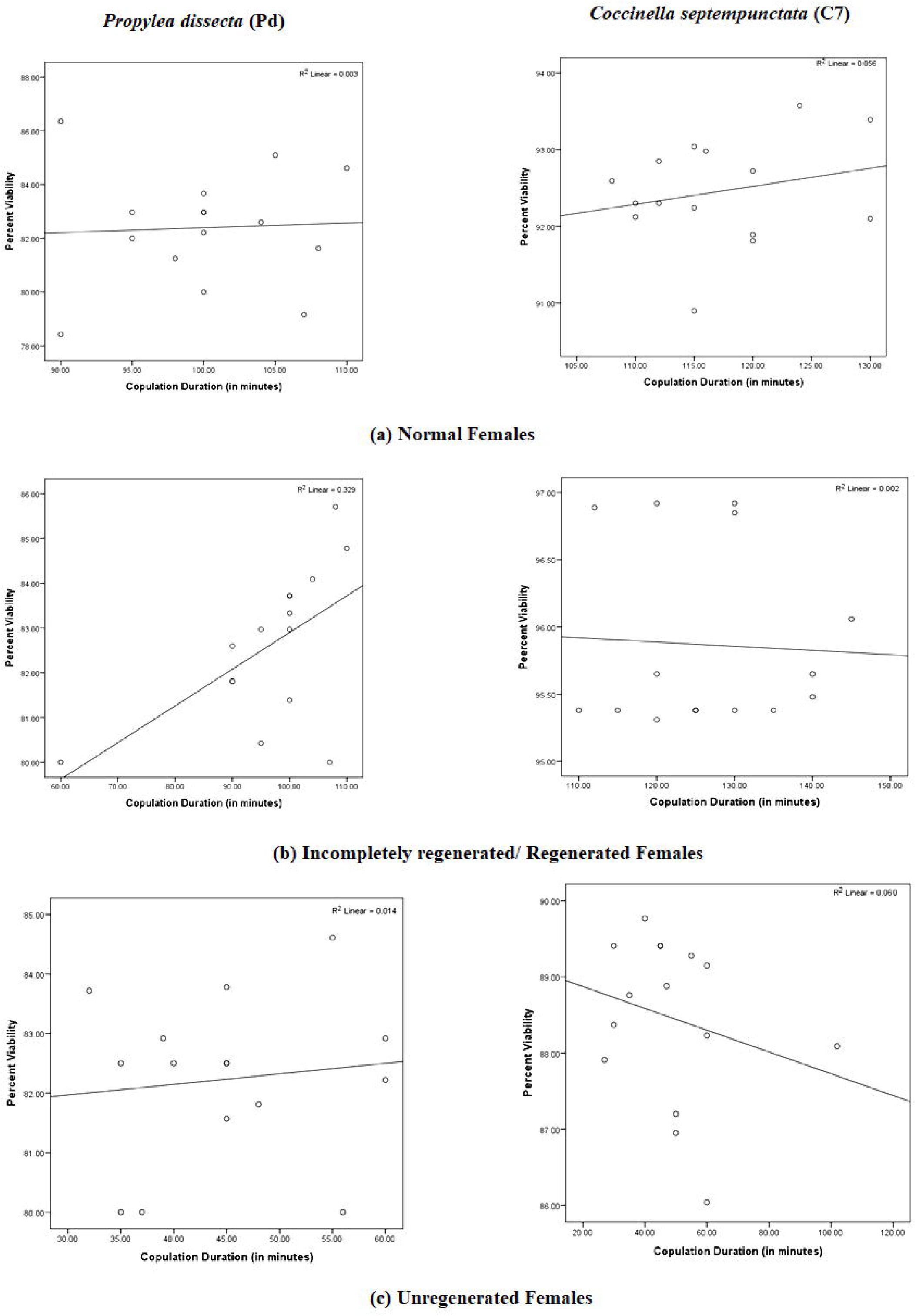
Relation between copulation duration and percent viability when different females of Pd and C7 mated with regenerated (incompletely in Pd) males.

**Figure 7:**
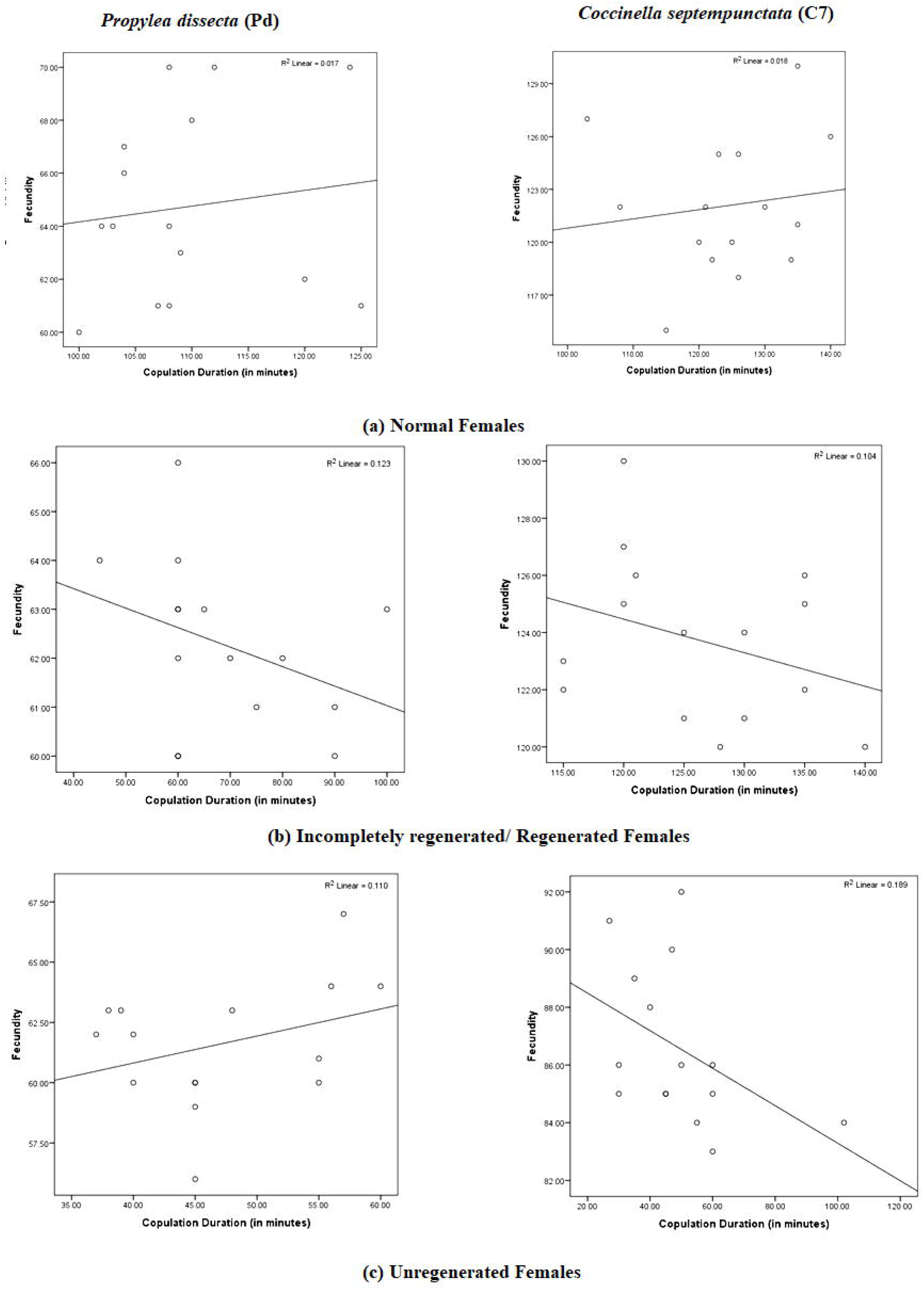
Relation between copulation duration and fecundity when different females of Pd and C7 mated with normal males.

**Figure 8:**
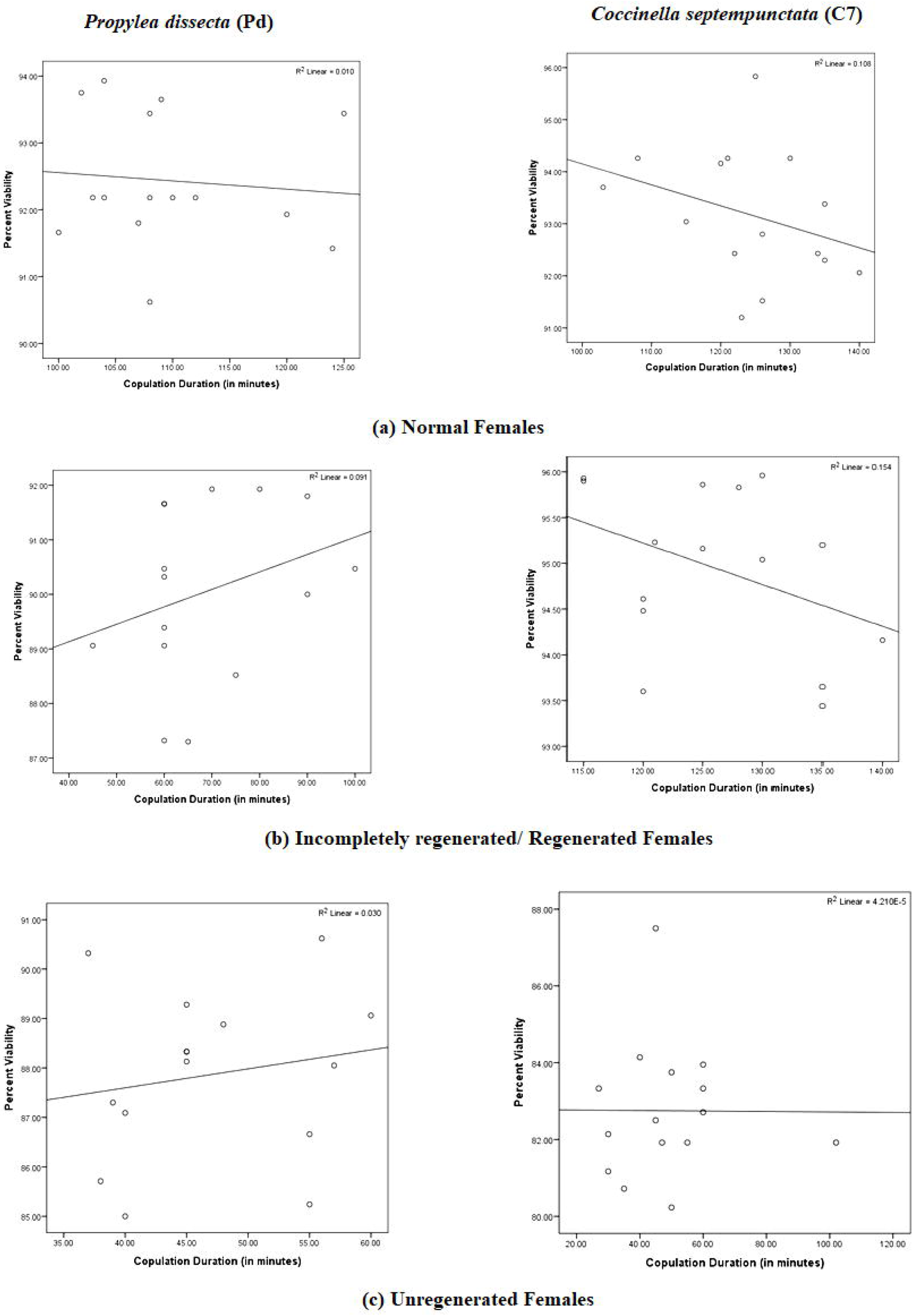
Relation between copulation duration and percent viability when different females of Pd and C7 mated with normal males.

**Figure 9:**
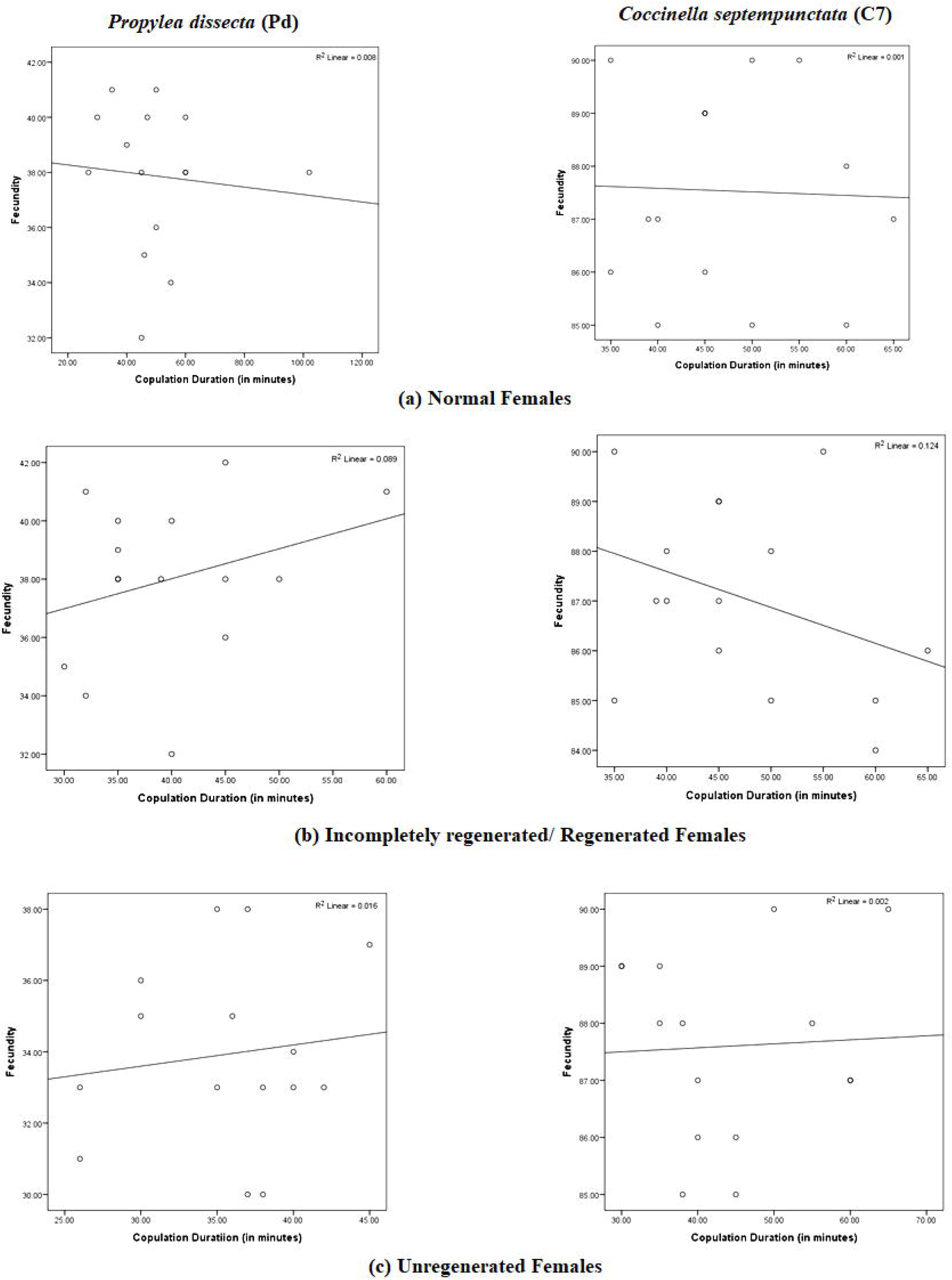
Relation between copulation duration and fecundity when different females of Pd and C7 mated with the unregenerated males.

**Figure 10:**
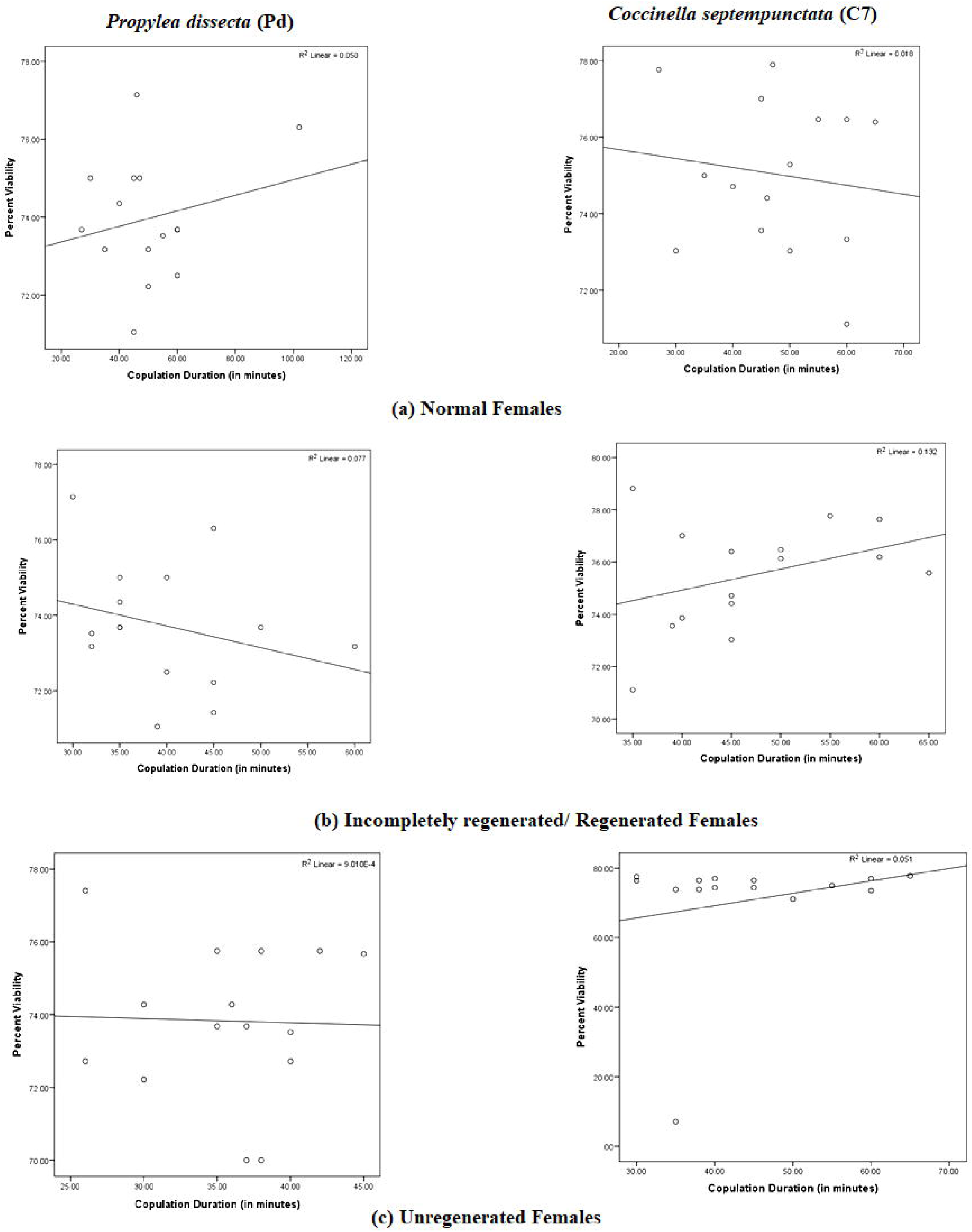
Relation between copulation duration and percent viability when different females of Pd and C7 mated with unregenerated males.

The regression analysis reveals that in *C. septempunctata* no positive relation were observed between copulation duration and reproductive parameters when different females were mated with regenerated, normal and unregenerated males (Fig.5,6,7,8,9,10). Similar results were also observed for the *P. dissecta*. There was no positive relation between the copulation duration and reproductive attributes in all the mating treatments (Fig.5,6,7,8,9,10).

## DISCUSSION

In the present study we found that longest time of commencement of mating (TCM) and shortest copulation duration were recorded for unregenerated treatments in both the ladybirds. Similarly, fecundity and percent egg viability were minimum in unregenerated treatments for both the ladybird species.

It was found that the unregenerated and incompletely regenerated adults (in *P. dissecta)* adults took more time to commence mating and mated for shorter duration as compared to regenerated and normal adults. This can probably be attributed to the rejection behaviour of female post assessment of the physical condition of males (Joseph *et al*.1999; Michaud 2003; Martini *et al*.2013). Another reason which could be explained for the lower performance of unregenerated and incomplete adults was incomplete physical contact with their mates due to the missing limb (Shandilya *et al*. 2018). This supports the hypothesis of the honest display of signals (Zahavi 1975; Grafen 1990 a,b; Zahavi & Zahavi 1997) where the individuals with more physical fitness get more chances for mating. Studies in insects and higher animals have revealed that the male ornamentation is inversely proportional to their fighting success and mating success (Moller 1990; Thornhill 1992; Swaddle & Cuthill 1994; Watson and Thornhill 1994; Hill 1995).

The reduced fecundity and percent egg viability by unregenerated and incomplete regenerated adults may be attributed to (a) the lower mating duration as reported by Haddrill *et al*. (2008) reported that longer copulation durations result in increased paternity share due to larger number of spermatophore transfer and (b) utilization of sperms by females owing to the perception of reduced fitness of males (Wang *et al*. 2015; Firman *et al*.2017). Differential usage of sperms by females owing to the status of the males has been well established (Kirkpatrick, 1987; Andersson, 1994). In some taxa, females are known to block the sperm of the unfit or unhealthy males (Edvardsson and Arnquivst, 2000, Kokko *et al*. 2003). Males with lower viability may also be attributed to the wrong positioning of the aedeagus due to lack of physical fitness. Study by Shandilya *et al*. (2018) revealed the reduced viability in leg impaired *M. sexmaculatus* can be due to the lack of structures which are required for proper holding of mates. No positive relation between the copulation durations and reproductive attributes were recorded in both the ladybirds in different mating treatments. This revealed that probably the assessment of the physical fitness by males and females may lead to the differential investment in mating leading to differences in fecundity and percent egg viability.

Thus, from the above study it can be concluded that regeneration in both the ladybirds modulates the mating and reproductive attributes and unregenerated adults are poor in mating and reproductive performances irrespective of the size of ladybirds.

## ACKNOWLEDGEMENTS

SS acknowledges Basic Science Research Fellowship by University Grants Commission, New Delhi, India (F.No.25-1/2014-15 (BSR)/7-109/2007/BSR) dated August 25, 2015.

